# Biological integrity enhances the qualitative effectiveness of conditional mice-oak mutualisms

**DOI:** 10.1101/2021.11.10.468095

**Authors:** Teresa Morán-López, Jesús Sánchez-Dávila, Ignasi Torre, Alvaro Navarro-Castilla, Isabel Barja, Mario Díaz

## Abstract

Scatter-hoarding decisions by rodents are key for the long-term maintenance of scattered tree populations. Decisions are determined by seed value, competition and predation risk, so that they can be influenced by the integrity of the biological system composed by trees, rodents, ungulate competitors, and rodent predators. We manipulate and model the oak-mice interaction in a Spanish dehesa, an anthropogenic savanna system suffering chronic tree regeneration failure, and quantify the joint effect of intrinsic and extrinsic factors on acorn dispersal effectiveness. First, we conducted a large-scale cafeteria field experiment, where we modified ungulate presence and predation risk, and followed mouse scatter-hoarding decisions under contrasting levels of moonlight and acorn availability. Then, we estimated the net effects of competition and risk by means of transition probability models that simulated mouse scatter-hoarding decisions according to the environmental context. Our results show that suboptimal conditions for mice balance the interaction towards the mutualism as they force mice to forage less efficiently. Under stressful conditions (predation risks and presence of ungulates), lack of antipredatory cover around dehesa trees limited transportation of acorns, but also precluded mice activities outside tree canopies. As a result, post-dispersal predation rates were reduced and large acorns had a higher probability to survive. Our work shows that inter-specific interactions preventing efficient foraging by scatter-hoarders benefitted seed dispersal. Therefore, the maintenance of the full set of producers, consumers, dispersers and predators in ecosystems is key for promoting seed dispersal effectiveness in conditional mutualisms.

## Introduction

Scatter-hoarders are key dispersers in temperate and Mediterranean forests where acorn-bearing trees (oaks) tend to be dominant [1-5]. Nut dispersal by scatter-hoarders (synzoochory) is a classical plant-animal conditional mutualism. The outcome of the interaction may be either mutualistic (dispersal) or antagonistic (predation) depending on whether seeds are consumed or, alternatively, cached and not retrieved [6]. The balance between mutualism and antagonism is contingent on intrinsic properties of interaction partners (e.g. propensity of animals to store food) as well as on the ecological setting in which the interaction occurs [5]. As a result, the net effects of synzoochory can be highly dynamic in space and time making difficult to predict its outcomes along environmental gradients and ecological timescales [7, 8].

Several mice species (*Apodemus, Mus, Peromyscus*) are the main scatter-hoarders in landscapes where avian dispersers (corvids; [9]) are absent, scarce or inefficient [2, 10]. Two main external factors modulate mouse scatter-hoarding decisions: competition for seeds and predation risks [11-13]. Intraspecific competition and the presence of ungulates tend to encourage seed mobilization [6, 14-16]. Especially, when predating seeds *in situ* is more time-consuming than storing them for later consumption [12] and shrubs provide enough antipredatory cover during transportation [16, 17]. Even though lack of antipredatory cover can limit dispersal [16], intermediate risks can promote mobilization when mice carry away seeds to manipulate them in safer locations [12].

Risk perception, in turn, depends on factors that affect exposure to predators (e.g. moonlight) and direct cues of their presence (e.g. scent) [19-24]. Overall, moderate level of stress for foraging mice (i.e. competition and predation risk) tend to unbalance the rodent-tree interaction towards its mutualistic side. In the absence of stress, rodents usually act as efficient seed predators consuming, immediately or soon afterwards, seed crops under the canopy of mother trees [5].

Beyond the environmental conditions of plant-animal encounters, seed size can affect the initial outcomes of the interaction (selected, eaten or cached) as well as post-dispersal processes such as germination and seedling survival. Larger seeds are usually selected and preferentially cached because they provide higher food rewards [3, 24-28]. In addition, seed size enhances post-dispersal seedling survival and establishment [29], which is a key component of dispersal effectiveness [30] in scatter-hoarder animals [3, 5, 31]. Nonetheless, the strength and even sign of acorn size effects on mouse foraging decisions are not unequivocal, but context-dependent. Larger acorns are most preferred when food is scarce [32-34], but may be avoided when longer handling times [27] diminish their profitability [35, 36] or result in unaffordable predation risks during manipulation [11, 12]. Therefore, a full picture of the location of mice in the antagonism-mutualism continuum [4] requires accounting for seed size effects on scatter-hoarding decisions as well as the influence of competition and risk.

In this context, dehesas represent an excellent study system to assess the main factors modulating mouse foraging decisions, and hence, dispersal. They are savanna-like habitats, simpler than natural forests but diverse enough to maintain all key elements influencing the oak-mice conditional mutualism. In spite of this, dehesas, as well as other man-made systems dominated by scattered trees, suffer from a chronic lack of tree regeneration that compromises its long-term sustainability [37, 38]. Depending on the local intensity of management, nearby areas can have contrasting levels of shrub cover, mice densities and competition with ungulates [39, 40]. In addition, the community of predators is simpler than in forest areas, facilitating the experimental manipulation of direct cues of risks [24]. In this work we take advantage of a large-scale experiment of ungulate exclosure in a Mediterranean dehesa to (1) quantify acorn size effects across different stages of the dispersal process (from seed choices to initial fates); and (2) evaluate if size effects are consistent across contrasting scenarios of predation risk and inter- and intraspecific competition. In addition, we parameterized a transition probability model that assembled all scatter-hoarding decisions by mice to quantify and tease apart the net effect of competition and risk on acorn dispersal. Our integrated approach combining field experiments and mechanistic modelling will allow testing whether the key role of rodents as seed dispersers in scattered tree systems can be enhanced by restoring the biological integrity of these systems [13].

## Methods

### Study area and species

Field work was carried out in the holm oak *Quercus ilex* dehesa woodlands of the Cabañeros National Park (Central Spain, Ciudad Real province, 39°24’ N, 38°35 W). Dehesas are savanna-like man-made habitats resulting from shrub removal and tree thinning and pruning to enhance herb growth for livestock [41]. The studied dehesas were opened in the late 1950s. Currently they have no livestock but wild ungulate populations of red deer *Cervus elaphus* and wild boars *Sus scrofa*. Deer densities were around 0.14 ind./ha [42] and boars are abundant but at unknown densities[43]. Acorns fall from trees from mid-October to late November [44].

The study area covers around 780 ha, with two ungulate exclosures (made with wire fences 2 m tall and 32 cm x 16 cm mesh) of 150 ha and 4.65 ha separated from each other by 1500 m. The exclosures prevent the entrance of ungulates but not of mesocarnivores (mainly common genets *Genetta genetta* and red foxes *Vulpes vulpes*; pers. obs. based on scat searches) and raptors. Both areas have similar tree abundance (average density 20.4 trees ha^-1^) and low shrub cover (<1%), as measured on aerial photographs and vegetation surveys both under canopies and outside them [24]. The Algerian mouse is the most abundant scatter-hoarding rodent in the area [45], and it is a common prey of genets and other generalist predators [46, 47].

### Experimental design

Tree occupancy by mice was established by means of live trapping using Sherman traps (23 × 7.5 × 9 cm; Sherman Co., Tallahassee, USA) baited with canned tuna in olive oil mixed with flour and a piece of apple. Water-repellent cotton was provided to prevent the cooling of the individual captured overnight. Traps were set during two consecutive days during the new moon of January 2012. High capture probability of *M. spretus* (detectability: 0.88±0.03SE; [48]) allowed to consider false negatives in occupancy unlikely. Among trees known to be occupied by Algerian mice, we randomly selected ten trees inside and ten outside in each of the two exclosures (40 focal trees in total).

We paired focal trees according to their proximity and we randomly assigned a predator scent treatment to one of them. Predator scent treatment consisted in placing fresh genet feces (10 g) mixed with distilled water close to a corner of the cages where acorns were placed [24]. Genets are generalist predators whose presence and scats are known to influence rodent behavior [17, 21, 23]. Fresh feces were collected from two captive common genets housed in the Cañada Real Open Center (Madrid, Spain).

Fresh acorns were collected from holm oaks growing near the study area in October 2011 and stored dry in a cooler (4 °C) until use. Sound acorns, with no marks of insect damage [45], were weighed with a digital balance to the nearest 0.01 g. Groups of 15 acorns in three categories were randomly selected (5 each, large, >10 g; medium-sized, 5-10 g, and small, 1-5 g). Acorns were placed under the canopy of each focal tree inside a 50 cm × 50 cm x 15 cm galvanized-steel cage to prevent acorn consumption by birds or ungulates [45]. A metal wire (ø 0.6 mm, 0.5 m length) with a numbered plastic tag was attached to each acorn [16]. After removing any naturally-present acorn within the cages, we randomly placed acorns in the intersection of a 3 rows x 5 columns grid. To track mouse choices, acorn size for each position was noted. Acorns were left exposed to mice for three consecutive nights, then removed. Mobilized acorns were located by looking at the plastic tags during the following days. To account for changes in night brightness and acorn availability [19, 22], the cafeteria experiment was repeated four times during the full-moon and new-moon periods of November 2011 and February 2012. No official permits or protocol approvals were legally necessary since we did not manipulate individual mice except for checking whether trees were occupied or not by means of live traps. We followed Guidelines of the American Society of Mammalogists for the use of wild mammals in research [49]. We performed all manipulations with disposable latex gloves, to avoid effects of human odor on rodent behavior [50].

### Mouse foraging behavior

A video-camera OmniVision CMOS 380 LTV (OmniVision, Santa Clara, USA) (3.6 mm lens) monitored mice foraging activity within each cage [17]. Cameras were set on 1.5 m tall tripods located 2.5 m from each cage, powered by car batteries (70 Ah, lead acid) connected to a solar panel (ono-silicon erial P_20; 20 w). Video-cameras were connected to ELRO recorders with dvr32cards (ELRO, Amsterdam, Netherlands) and took continuous record for three consecutive days autonomously (recorded in quality at 5 frames s^−1^). Events with rodent activity, from the entry of the individual into the cage up to the exit from it, were located and separated using Boilsoft Video Splitter software (https://www.boilsoft.com/videosplitter/)[17]. Within each event we noted which acorn was manipulated and whether it was removed outside the cage. For removed acorns we measured mobilization distance (cm) and noted its status (predated or not after transportation).

### Data analysis

To assess acorn choice by rodents, we fitted a hierarchical multinomial model. Our response variable was acorn selection (yes/no). Our explanatory variables were: acorn size (g), moon phase (new/full), month (February, November), ungulate presence (yes/no), predator scent (yes/no), acorn availability in the cage (g) and the two-way interactions between size and environmental effects. Local acorn availability was measured as total acorn mass in the cage during the event. Both, acorn size and availability were scaled previous to the analyses. Focal tree was introduced as a random factor in the intercept term. To evaluate the effects of acorn size on the probability of removal, we used a hierarchical logistic model. Our response variable was whether a selected acorn was mobilized outside the cage or not (yes/no). Our explanatory variables and random effects were the same as in the multinomial model.

Subsequently, we analyzed the effect of acorn size and environmental covariates (and their two-way interaction) on seed dispersal. Our response variables were mobilization distances (cm, log-transformed) and deposition status (viable or predated). We used a hierarchical Gaussian model in the former case, and a hierarchical logistic model in the latter. Our explanatory variables and random effects were the same as in the previous models. In all four models (selection, removal, mobilization distance and fate) we used uninformative priors (Supplementary File 1). All analyses were performed employing a Bayesian approach with JAGS 3.4.0 [51]. We checked for convergence for all model parameters (Rhat < 1.1) and that the effective sample size of posterior distributions was high (>800). We estimated mean and credible interval of posterior distributions, calculated the proportion of the posterior distribution with same sign of the mean (f) and evaluated the predictive power of our models by means of posterior predictive checks (Supplementary Files 1 and 2).

### Simulating scatter-hoarding decisions

To estimate the joint effect of seed size, competition and risk on acorn dispersal we designed a probability transition model in which simulated mice adapted their foraging behavior to the environmental context (Supplementary File 3). Before model run, we parameterized mouse scatter-hoarding decisions (from selection to initial fate) following the same scheme of regressions explained in the previous section. Nonetheless, here we only used data from November, the period of peak acorn falling in our study system.

Consequently, we did not include month as a covariate. For each behavioral submodel (selection, removal and initial fate), we obtained posterior distributions of parameters by running 50000 iterations in three chains (in all cases Rhat< 1.1, and Neff> 1000).

Model setup mimics our experimental design, 20 trees outside and 20 inside exclosures paired according to a predator scent treatment (presence vs. absence). Simulations begin under new moon conditions with focal trees offering 15 acorns of large, medium and small sizes (5 each). Acorn size is sampled from empirical distributions of these size categories. In each focal tree, the number of foraging events is drawn from a Poison distribution with mean equal to the average number of events observed in the corresponding moon phase 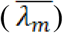. During each foraging event, simulated mice decide which acorn to handle and whether to remove it or not. If removed, mice decide to predate it or not after mobilization and acorn availability in the cage is updated. Once all foraging events (of all trees) are simulated, acorn dispersal is modelled under full moon conditions (Supplementary File 3, Fig. S1).

For each model run we sampled parameter of behavioral submodels (selection, removal and deposition) from posterior distributions fitted to data (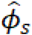, being s each behavioral submodel and *Φ* its parameter set). Thus, in our simulations, mice adapted their decisions to acorn size and availability (in the experimental cage), characteristics of the focal tree (i.e. ungulate and predator scent presence), and the moon phase in which the foraging event occurs (new or full moon). After each model run (dispersal under new and full moon conditions), the program tracked the size and status of handled acorns and the environmental covariates in which the foraging event occurred. We run the model 1000 times and plotted deposition rates of viable acorns and their size with respect to the moon phase and tree characteristics (predator scent and ungulate presence). See Supplementary File 3 for detailed model specifications and Fig. S1 for a summary of the process overview.

## Results

Before setting the cafeteria experiments in November, we removed from cages 53.3 acorns/m^2^ on average (range: 0-104). No acorn was found in February. We monitored 2280 acorns under 38 focal trees. We detected mouse activity in 18 and 26 trees in the new and full moon of November, and in 26 and 24 trees in the new and full moon of February, respectively. Mice manipulated 1378 acorns. Out of them, 505 were mobilized outside cages and 385 (76%) were relocated [26].

### Foraging decisions in the focal tree: selection and removal

In general, mice selected larger acorns, but the positive effect of size was modulated by environmental conditions. Size-driven selection preferentially occurred in the absence of competition with ungulates (Fig. 1A) and predator scent (Fig. 1B). In addition, mouse selectivity was enhanced under low local acorn availability (Table 1, selection). Among selected acorns, mice preferentially removed smaller ones. Such selective behavior occurred when risks were low due to reduced night brightness (new moon, Fig. 1C) or lack of predator scent (Fig. 1D), as well as when ungulates were absent (Table 2). Acorn availability at local and landscape scales did not modify size effects, although they changed mobilization rates. Rates were higher in lean periods (13% in November *vs* 24 % in February) and when local availability was high (Table 2, removal).

**Fig. 1.**
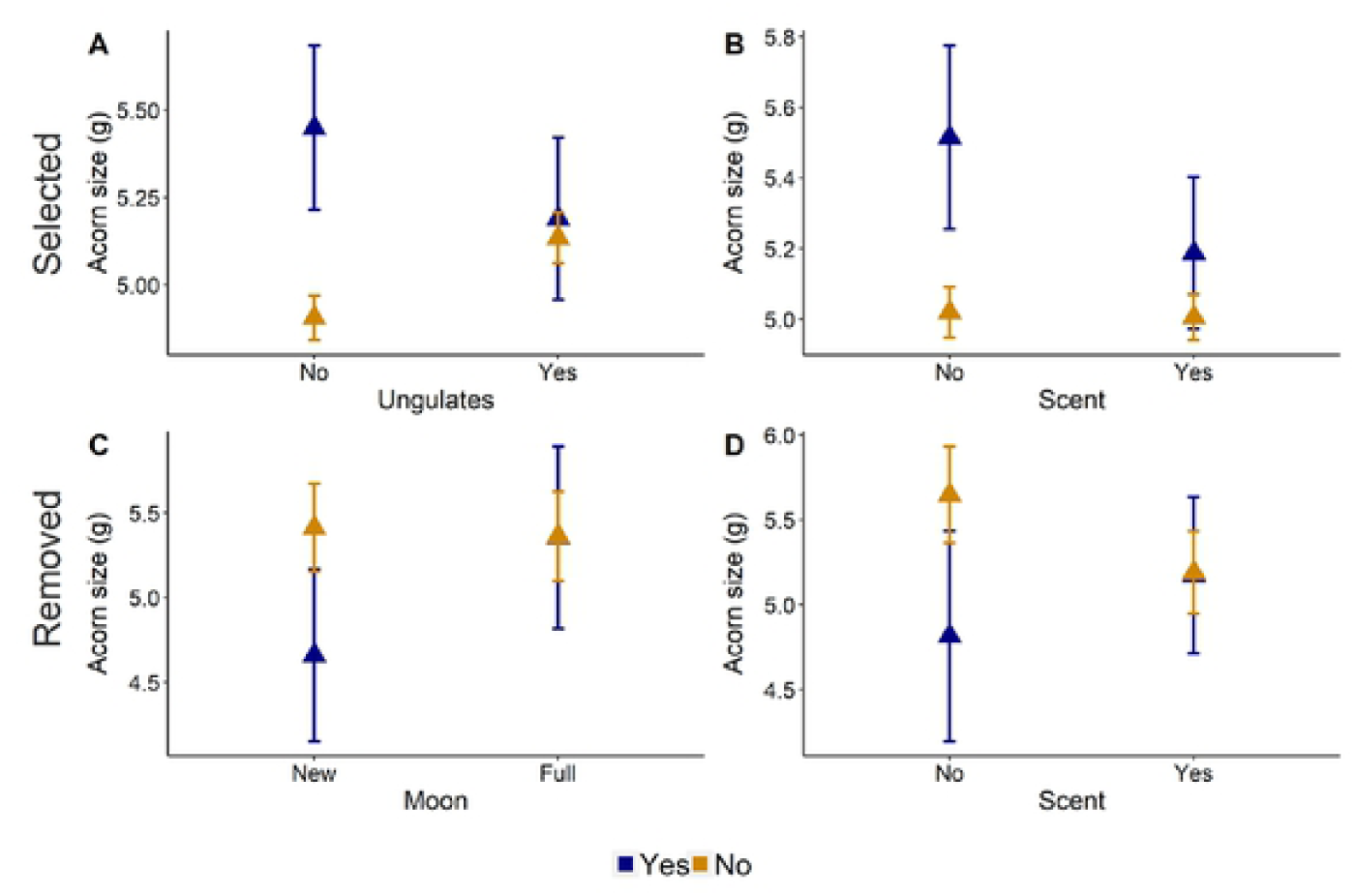
Mouse foraging decisions during acorn selection and removal (upper and lower panels, respectively). Mouse preferentially selected larger acorns. However, such selective behavior only occurred when (A) ungulates were absent and (B) there was no predator scent. Among selected acorns, mouse tended to mobilize smaller ones. This selective behavior occurred when risks were low due to (C) new moon conditions or (D) lack or predator scent. Also, when ungulates were absent (Table 1). Points represent mean values, bars standard errors.

**Table 1.** Summary table of the effects of size, moonlight, month, ungulate presence, predator scent and local acorn availability (and their interactions with size) on the probability of acorn selection and removal.

| Process | Fixed effect | Mean | HPD1 | f |  |
| --- | --- | --- | --- | --- | --- |
| Acorn selection | **Size** | **0.19** | **[0.09, 0.29]** | **1.00** | **\*\*** |
|  | Moon (Full) | 0.03 | [-6.22, 6.35] | 0.50 |  |
|  | Month (February) | -0.03 | [-6.12, 6.06] | 0.50 |  |
|  | Ungulate (Yes) | -0.05 | [-6.27, 6.16] | 0.51 |  |
|  | Scent (Yes) | 0.01 | [-6.16, 6.28] | 0.50 |  |
|  | Availability | -0.02 | [-6.32, 6.19] | 0.50 |  |
|  | Size*Moon | 0.07 | [-0.03, 0.17] | 0.93 |  |
|  | Size*Month | -0.06 | [-0.16, 0.04] | 0.88 |  |
|  | **Size\*Ungulates** | **-0.13** | **[-0.23, -0.03]** | **0.99** | **\*\*** |
|  | **Size\*Scent** | **-0.08** | **[-0.18, 0.01]** | **0.96** | **\*** |
|  | **Size\*Availabilitiy** | **-0.04** | **[-0.09, 0.01]** | **0.95** | **\*** |
| Acorn removal | **Size** | **-0.50** | **[-0.94, -0.07]** | **0.99** | **\*\*** |
|  | Moon (Full) | 0.07 | [-0.27, 0.39] | 0.65 |  |
|  | **Month (February)** | **0.77** | **[0.43, 1.11]** | **1.00** | **\*\*** |
|  | Ungulate (Yes) | -0.22 | [-0.96, 0.47] | 0.73 |  |
|  | Scent (Yes) | 0.20 | [-0.53, 0.93] | 0.72 |  |
|  | **Availability** | **0.29** | **[0.12, 0.46]** | **1.00** | **\*\*** |
|  | **Size\*Moon** | **0.29** | **[-0.02, 0.60]** | **0.96** | **\*** |
|  | Size*Month | -0.09 | [-0.41, 0.22] | 0.71 |  |
|  | **Size\*Ungulates** | **0.24** | **[-0.07, 0.55]** | **0.94** | **•** |
|  | **Size\*Scent** | **0.30** | **[0.00, 0.59]** | **0.98** | **\*** |
|  | Size*Availabilitiy | 0.10 | [-0.07, 0.26] | 0.88 |  |

Mean of posterior distribution, highest posterior density interval (HPD) and percentage of the posterior
distribution with the same sign as the mean (f) are shown. Effects with $f \geq 0.95$ are in bold. • depicts $f \in$

**Table 2.**
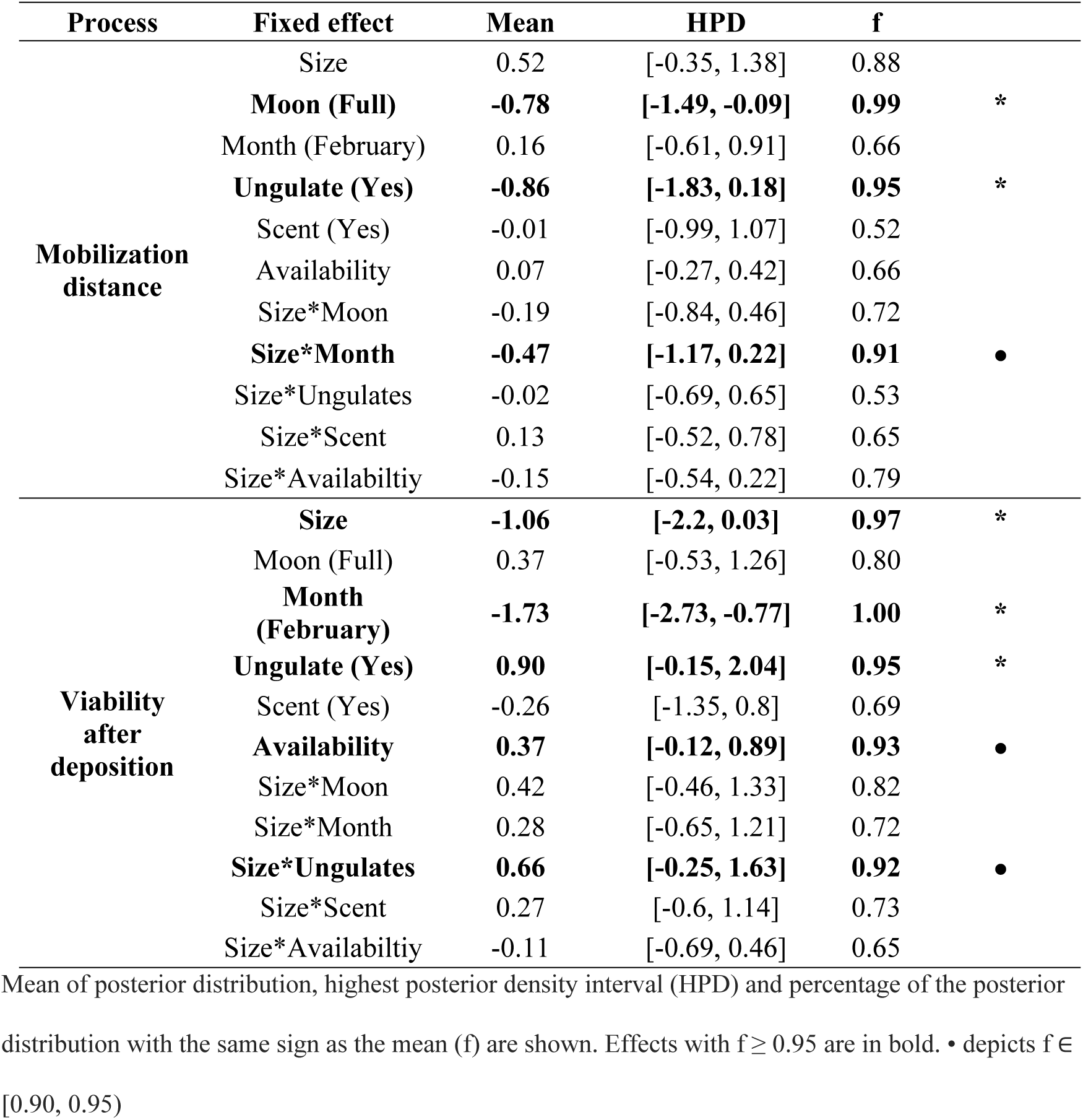
Summary table of the effects of size, moonlight, month, ungulate presence, predator scent and local acorn availability (and their interactions with size) on acorn mobilization distances and the probability that it is deposited in a viable status (vs predated).

### Foraging decisions during dispersal: mobilization distances and predation

Mice mobilized acorns closer under new moon conditions (Fig. 2A) and when ungulates were present (Fig. 2B). Even though acorn size did not affect overall mobilization distances, in February mice tended to mobilize smaller acorns further away (Table 2, Mobilization distances: Size*Month). Regarding initial fates, post-dispersal predation increased during lean periods (February) and was relaxed in the presence of ungulates (Table 2, Viability after deposition). In addition, larger acorns were preferentially consumed (Fig. 2C), though the presence of ungulates attenuated this negative effect (Fig. 2D).

**Fig. 2.**
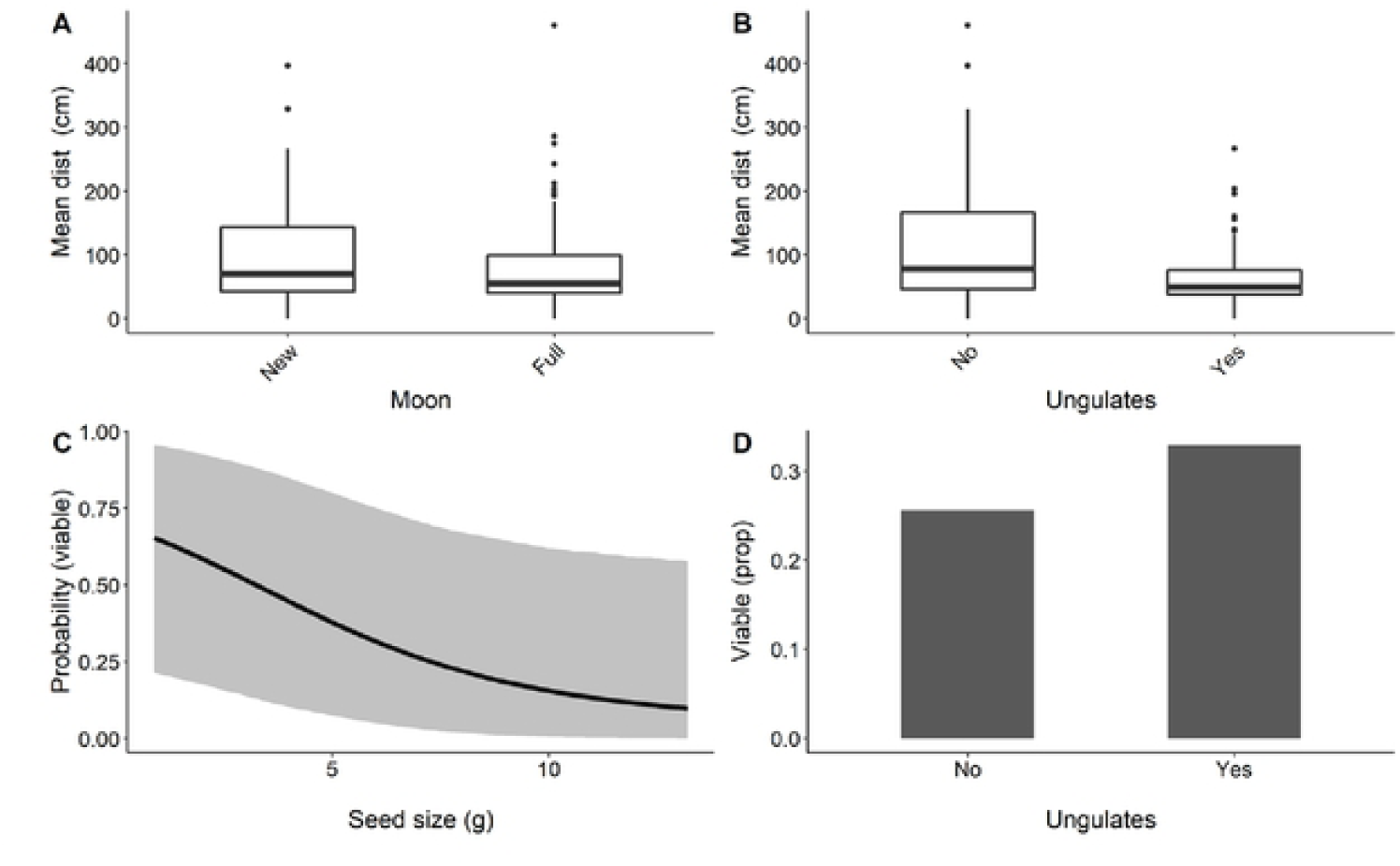
Summary of mouse foraging decisions during and after mobilization. (A) Under full moon conditions and (B) in the presence of ungulates mice mobilized acorns closer. Regarding deposition, (C) larger acorns had a lower probability of escaping predation as well as (B) those mobilized in areas with ungulates. Black line in panel C represents mean effects of acorn size on the probability of escaping predation, and shaded area 0.95 credible intervals. Panel D, represents the proportion of viable acorns (after deposition) from trees inside and outside ungulate exclosures.

### Transition probability model for acorn dispersal

Under optimal conditions (new moon, no predator scent or ungulates), post-dispersal predation rates were higher (Fig. 3A) and simulated mice preferentially consumed large acorns (i.e. viable acorns were smaller, Fig. 3B-D, left bars). However, predation risks and ungulate presence precluded acorn consumption after mobilization and attenuated selection. As a result, the proportion of viable acorns increased and they were larger (Fig. 3A and B-D, right bars).

**Fig. 3.**
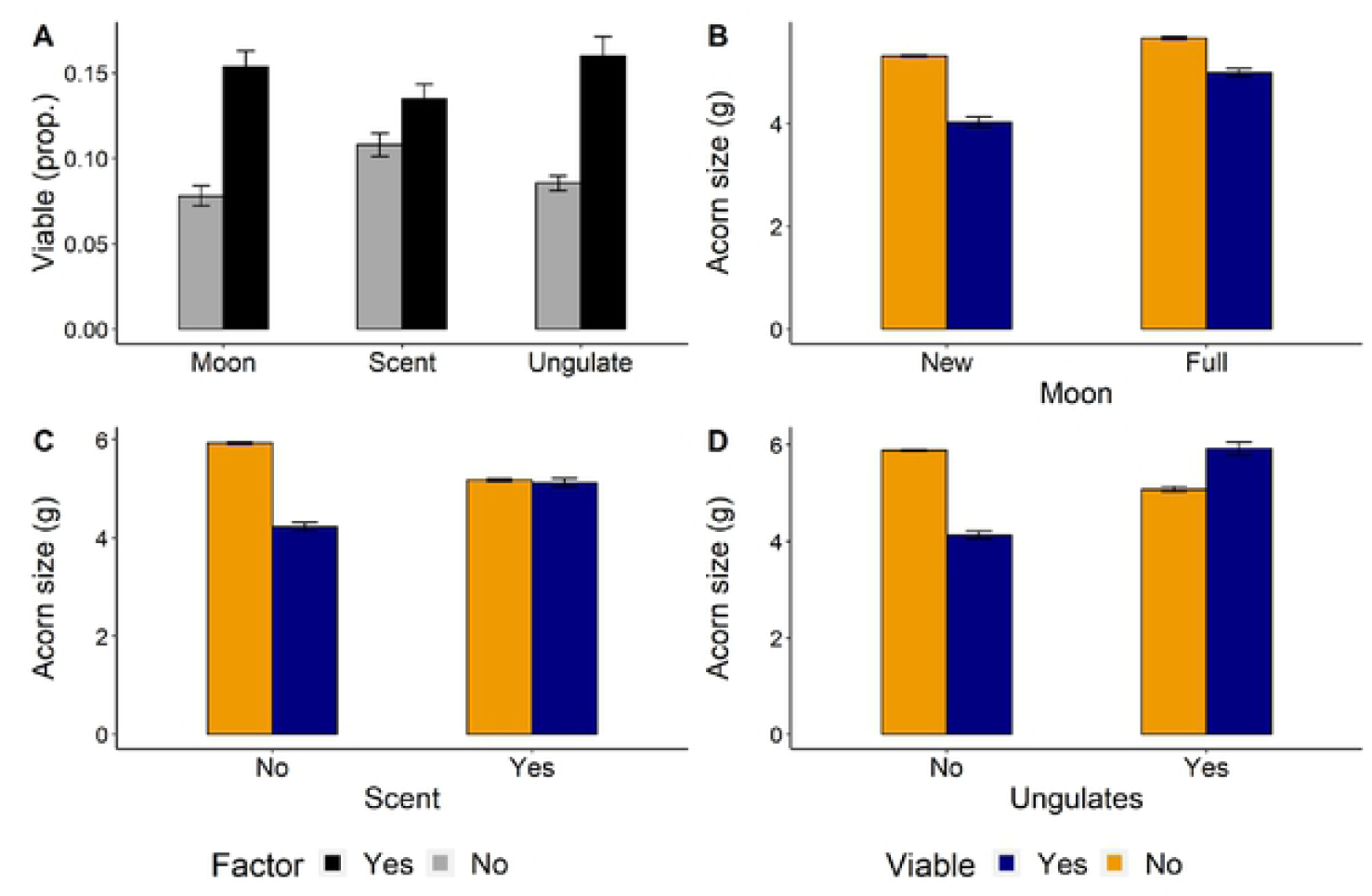
Results from simulations of the probability transition model for acorn dispersal. (A) Under more stressful conditions (black bars), the proportion of acorns escaping predation increased. In general, mice tended to predate larger acorns under (B) new moon conditions, (C) in the presence of predator scent and (D) when competing with ungulates. Under more stressful conditions (Fig. B-D, right bars), mice were less selective and consequently size effects on acorn predation were attenuated (B), disappeared (C) or reversed (D). Bars represent mean values (±s.e.) across 1000 simulations.

## Discussion

Overall, our results show that environmental stress unbalances the oak-rodent interaction towards the mutualism side. When relaxed, mice preferentially consumed large acorns and removed small ones. Furthermore, mobilized seeds were more likely to be predated. In contrast, under stressful conditions (predation risk and competition) mice foraged opportunistically and reduced their activity outside tree canopies. As a result, predation rates after mobilization decreased, and larger acorns had a higher probability to survive. This bolsters the idea that interactions with third-party players can strongly affect seed dispersal effectiveness of scatter-hoarders [12, 15, 52]. Also, that intermediate stress can benefit plants by reducing the capacity of scatter-hoarders to recover mobilized seeds.

As expected, larger and more valuable acorns were preferentially handled by mice, which adapted this behavior to the environmental context [12]. In line with previous work, mice foraged opportunistically in trees with predator scent, probably because they devoted more time to vigilant behaviors [17, 27] at expenses of acorn discrimination [22]. In contrast, acorn availability effects did not follow the expectations of increased selectivity in scenarios of food depletion [28, 52, 53]. Seed size effects were similar between acorn fall peaks and lean periods. Furthermore, mice foraged randomly when ungulates were present, while they selected larger seeds within exclosures. These unexpected results can be explained by some particularities of our system. Dehesas are characterized by high acorn production and scarce shrub cover (<1%) around trees [41, 54, 55]. Under such circumstances, the effects of increased predation risks outside tree canopies can outweigh those of competition leading to a rapid and random harvesting of seeds [22]. In contrast, within ungulate exclosures, reduced grazing and soil compaction has promoted taller resprouts under canopies and increased cover of herbs and tussocks around trees [56]. As a result, in the absence of ungulates mice can forage under shelter and select the most profitable food items [17]. Taken together, these patterns point out that in dehesas, risk rather than competition modulates the effects of ungulate presence on acorn selection.

Larger acorns tend to be carried away, mobilized farther and preferentially cached in forests habitats [4, 26, 27, 34]. However, in our study larger acorns had a higher probability of being predated (*in situ* and after transportation) and seed size did not affect mobilization distances. Again, these results highlight that environmental conditions in dehesas are particularly harsh for rodents. In general, small seeds are preferentially mobilized when the costs of carrying large ones result unaffordable [12]. In the presence of ungulates, low antipredatory cover and high trampling risks may have triggered transportation costs [13, 57], deterring mice from carrying large seeds away.

Again, seed size effects were not fixed, but depended on direct and indirect cues of risk. Preferential removal of small seeds only occurred in trees with no predator scent or under new moon conditions, reflecting that only when risks are mild mice can take the time to select among the seeds available [15, 22].

Regarding post-dispersal survival, we expected higher predation when acorns were deposited close to tree canopies [13, 16]. Nonetheless, this relationship blurred in our system. Outside ungulate exclosures, larger acorns had a higher probability of escaping predation in spite of being mobilized nearby source trees. In dehesas, the pervasiveness of open land cover forces mice to concentrate their activities beneath canopies [13, 24, 56], and decreases the likelihood that mobilized acorns are encountered and consumed [58]. Taken together, our results suggest that intermediate levels of stress can enhance seed dispersal effectiveness by mice (as suggested by [52, 59]). Accordingly, in our simulations, suboptimal conditions (due to increased risks or ungulate presence) enhanced dispersal. Increased risks discouraged mice from investing time in selecting which acorns to carry away and from consuming seeds after mobilization.

Consequently, predation rates were reduced and larger acorns had a higher probability of dispersal. In Mediterranean systems, seedlings from larger acorns are more resistant to summer drought [29, 60], which represents the main recruitment bottleneck for oak regeneration [38, 61]. Thus, our simulation results suggest that suboptimal conditions can enhance both, the quantity and quality component of dispersal effectiveness by mice.

This work builds on previous research analyzing the effects of competition and risk on mouse foraging behavior in dehesas [17]. Here, by accounting for all stages of scatter-hoarding decisions (from initial manipulation to consumption after mobilization [4], and including the entire acorn fall season [27] as well as contrasting moon light conditions [22], we obtained a more in-depth understanding of the main drivers of dispersal effectiveness. Moreover, our transition probability model allowed us to assemble all stages of the scatter-hoarding process, and hence, to estimate the net effects of competition and risks on initial seed fate. However, future work that analyzes the actual probability of recruitment of mobilized seeds is needed. In the Mediterranean area, seedling recruitment usually concentrates under the shade of shrubs, where conditions are milder [62-64]. Thus, it remains an open question whether higher rates of seedling dry out in dehesas (due to lack of shrub cover) can outweigh the benefits provided by enhanced cache survival. Once information about long-term survival of caches and seedling recruitment is available, it can be easily included in our model [65].

## Concluding remarks

Our mechanistic approach provides new insights about the joint effect of habitat structure, competition and risk on dispersal effectiveness in synzoochorus interactions. In particular, we show that suboptimal conditions for scatter-hoarders can balance the interaction towards the mutualistic side. High predation risks forced mice to forage less efficiently resulting in a higher probability of post-dispersal survival of large acorns.

Our work points out that environmental stress can be an important factor modulating the spatial and temporal dynamism of synzoochorous interactions [5]. Also, it supports the view that biological integrity (presence of the full set of producers, consumers, dispersers and predators) is key for ensuring seed dispersal effectiveness in synzoochorous conditional mutualisms [5, 52]. This may be particularly important in man-made habitats like dehesas, which depend on conditional mutualisms to ensure their long-term sustainability [38, 55].

## Supporting information

**S1. Structure of models and priors**

**S2. Posterior predictive checks**

**S3. Specifications of transition probability model for acorn dispersal S4. Code for the transition probability model**

**S5. Databases**

## Acknowledgements

D. López, M. Fernández and C. L. Alonso helped during fieldwork. D. López, B. Ramos and M. de Pablo pre-processed the video recordings, and D. Gallego, D. Valero, A. Velasco, C. J. González and E. Sánchez visualized the recordings noting seed choices. Authorities of the Cabañeros National Park provided the official permissions to carry out field experiments. J. España provided common genet scats. This study is a contribution to the projects RISKDISP (CGL2009-08430) and VULGLO (CGL2010-22180-C03-03), funded by the Spanish Ministry of Economy, and REMEDINAL3-CM (S2013/MAE-2719), funded by the Autonomous Community of Madrid. We declare no conflict of interest.

## Author contributions

MD conceived and executed the field experiment with the aid of IT, IB, TM-L and AN-C. JS-D compiled the data and performed preliminary analyses and drafts. TM-L executed the final data analyses and proposed the final main focus of the paper. TM-L and MD wrote the final version of the paper on former versions drafted by JS-D and contributed by all authors.

